# MFAP5 Drives Elastic Fiber Disorganization to Promote Pulmonary Fibrosis

**DOI:** 10.64898/2025.12.09.693350

**Authors:** Zhen Chen, Fanhao Kong, Lixin Huang, Wenze Wu, Ziping Wang, Hanbin Chen, Junting Zhang, Zixin Liu, Jin-Song Bian, Xiaowei Nie

## Abstract

**BACKGROUND:** Elastic fibers (EF) disorganization contributes to increased tissue stiffness and impaired lung function in pulmonary fibrosis (PF). However, the complex structural features of EF are difficult to capture using conventional histology. Moreover, the molecular mechanisms governing EF homeostasis during fibrosis remain poorly understood.

**METHODS:** A high-dimensional Elastic Fiber Algorithm (EFA) was developed to digitally quantify EF structural features in PF. Gene variation-rate analysis was performed to identify candidate regulators of EF homeostasis. Fibroblast-specific and pathological fibroblast-specific MFAP5 knockout mice were generated to assess the role of MFAP5 in PF rodent models. Bulk RNA sequencing was employed to investigate MFAP5-mediated fibroblast activation and signaling pathways.

**RESULTS:** EFA revealed profound EF disorganization in fibrotic lungs, capturing various architectural alterations. Gene variation-rate analysis identified MFAP5 as a candidate regulator of EF homeostasis, with upregulated expression in PF patients and mouse models. Fibroblast-specific deletion of MFAP5 significantly attenuated fibrosis, restored EF architecture, reduced collagen deposition, and improved pulmonary function in PF mouse models. MFAP5-positive fibroblasts displayed a dynamic shift toward pathological states during PF progression. Mechanistically, MFAP5 promoted fibroblast activation and ECM production via αvβ3 integrin-mediated TGFβ signaling.

**CONCLUSIONS:** MFAP5 is a key orchestrator of EF remodeling and fibroblast activation in PF. Targeting MFAP5 restores EF homeostasis, reduces fibrotic severity, and represents a potential therapeutic strategy for PF.

## INTRODUCTION

Elastic fibers (EF) are essential components of the lung extracellular matrix (ECM) that provide elasticity and structural integrity during respiration^1^. They are mainly composed of elastin and microfibrils, which together form a durable and highly organized network^1^. In fibrotic lung diseases, such as idiopathic pulmonary fibrosis (IPF) and bleomycin (BLM)-induced experimental fibrosis, abnormal elastin deposition and disorganized EF assembly are frequently observed^2–4^. These changes contribute to increased tissue stiffness, impaired lung compliance, and irreversible loss of function. Despite growing recognition of their importance, the molecular mechanisms that regulate EF homeostasis during fibrosis remain poorly understood. Additionally, current histological or imaging approaches largely rely on conventional staining or qualitative assessment, which fail to capture the complex structural and spatial features of EF^4^. As a result, the dynamic remodeling of EF during fibrosis has not been quantitatively or systematically characterized.

Microfibril-associated glycoprotein 5 (MFAP5), an extracellular matrix protein localized within microfibril-rich structures, is known to interact with components such as fibrillin and fibulin^5^. MFAP5 has been implicated in tissue remodeling, fibrosis, and cancer progression, where it can modulate fibroblast activity and extracellular matrix organization^6^. Given that MFAP5 is positioned at the interface between microfibrils and surrounding matrix proteins, it may act as a key regulator linking microfibril structure to signaling pathways that control fibroblast behavior. Emerging evidence suggests that MFAP5 expression is upregulated in fibrotic tissues^7–9^, but whether it contributes directly to EF dysregulation in pulmonary fibrosis (PF) remains unclear.

In this study, we developed an Elastic Fiber Algorithm (EFA) to assess EF characteristics in PF. Through gene variation-rate analysis, MFAP5 emerged as a central candidate regulating EF homeostasis during PF. Using BLM- or silica-induced PF mouse models, we demonstrated that fibroblast-specific deletion of MFAP5 ameliorated PF. Mechanistically, MFAP5 promoted fibroblast activation through αvβ3 integrin-mediated signaling, leading to disorganized EF assembly and exacerbated ECM deposition. These findings identify MFAP5 as a key molecular orchestrator of EF remodeling in fibrotic lung disease.

### Study Approval

All human samples were obtained from the Shenzhen People’s Hospital (China) and Wuxi People’s Hospital (China) with the informed consent of the patients, which was approved by the Ethics Committee for the Use of Human Subjects following The Code of Ethics of the Helsinki Declaration of the World Medical Association. All animal studies followed the guidelines of the Committee on Animal Research and Ethics of the Southern University of Science and Technology (China).

### Data Availability Statement

Source data and images for this study are available from the corresponding author upon reasonable request.

## RESULTS

### Multimodal Characterization of Elastic Fiber in PF

To capture the spatiotemporal dynamics of EF characteristics during PF, we developed a high-dimensional EFA for analysis (**Figure 1A**). EFA enables the digitalization of abstract EF features, facilitating the observation and quantification of EF alterations. Victoria blue/fast red-stained images were obtained from lung tissues of bleomycin (BLM)-treated mice at 21 days post injury (dpi), representing EF (blue) and cell nuclei (red) (**Figure 1B**). By applying EFA, Victoria blue/ fast red-stained images are converted into digital EF binary architecture images (**Figure 1C**). Nuclei were removed to prevent them from confounding EFA analysis, while the abnormal EF architecture remained clearly remained. Each image was quantified for more than 100 architectural features, capturing both local (such as fiber thickness and continuity) and global (such as network anisotropy and density heterogeneity) structural characteristics. UMAP visualization of the digitalized EF features from BLM-treated mouse lungs showed a left-lower shift compared with saline-treated controls (**Figure 1D**). These results demonstrate that EFA effectively captures fibrosis-associated alterations in EF architecture, providing a robust framework for high-dimensional structural analysis during pulmonary fibrosis.

**Figure 1.**
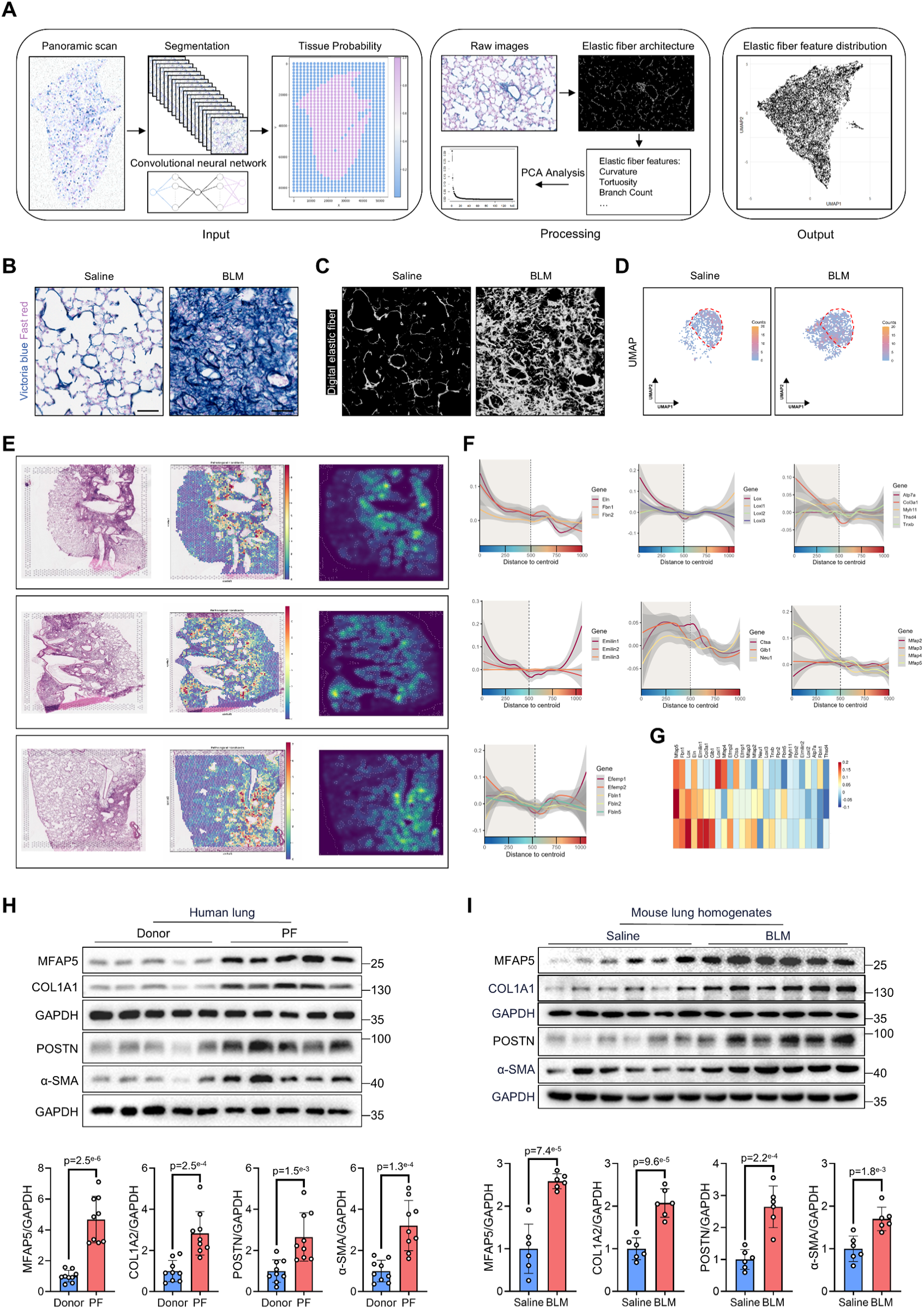
Elastic Fiber Algorithm-based structural profiling of elastic fibers in pulmonary fibrosis. **A**, Workflow of the Elastic Fiber Algorithm (EFA) for high-dimensional analysis of elastic fibers (EF) characteristics. EFA consists of three major components. Input module performs image tiling and tissue-background classification; Processing module separates the EF channel, reconstructs the digital EF architecture, and extracts high-dimensional structural features. Output module generates UMAP-based visualizations to summarize EF feature patterns. **B**, Representative Victoria blue/fast red-stained lung sections from saline- or bleomycin (BLM)-treated mice at 21 days post-injury (dpi). EF (blue) and nuclei (red). Scale bars=100 μm. **C**, Conversion of histological images into digital EF binary architecture images using EFA. **D**, UMAP visualization of digitalized EF features, revealing a characteristic shift in BLM-treated mouse lungs compared with controls. **E**, Identification of fibrotic hotspots by mapping scRNA-seq data onto spatial transcriptomics using cell2location, based on the spatial probability of *Postn⁺Cthrc1⁺* fibroblasts. **F**, Gene variation-rate analysis of EF-related genes within fibrotic hotspots. **G**, Integrative ranking of variation rates across 26 EF-related genes identifies MFAP5 as a candidate with consistently high variation rates across samples. **H**, Increased MFAP5 expression in lung tissues from patients with pulmonary fibrosis (PF), accompanied by elevated COL1A1, POSTN, and α-SMA expression. **I**, Upregulation of MFAP5 and fibrotic markers in BLM-induced PF mouse lungs. Data represent the mean±SD. Comparisons of parameters were performed with an unpaired Student’s t-test, followed by Tukey honestly significant difference test for multiple comparisons. COL1A1, collagen type I alpha 1 chain; POSTN, periostin; and α-SMA, alpha-smooth muscle actin.

### Elastic Fiber-Related Genes Dynamics in PF

To identify genes potentially involved in EF alterations in PF, an EF-related gene cluster was curated from the STRING, BioGRID, and Gene Ontology databases. This cluster encompasses 26 genes across seven categories, including ELN, FBN1, FBN2, and others. By mapping single-cell RNA sequencing data (GSE207851)^10^ onto the spatial transcriptomics dataset (S-BSST1409)^11^, fibrotic hotspot regions were defined based on the spatial probability of *Postn⁺Cthrc1⁺* fibroblasts (**Figure 1E**). Then, we conducted gene variation-rate analysis on the EF-related genes within the fibrotic hotspots. Results showed that genes involved in the ELN synthetic pathway (such as ELN and GLB1) exhibited high variation rates, suggesting that elastin contributes to the abnormal structural characteristics of EF within the fibrotic hotspots (**Figure 1F**). Similarly, other EF components, including FBN1 and FBN2, also displayed high variation rates (**Figure 1F**). Integrative analysis of variation rates across all EF-related genes identified MFAP5 as a candidate gene. MFAP5 exhibited consistently high variation rates across all samples (**Figure 1G**), implying its potential role in regulating EF homeostasis in PF. Importantly, MFAP5 was markedly upregulated in lung tissues from patients with PF, accompanied by increased expression of fibrotic markers such as COL1A1, POSTN, and α-SMA (**Figure 1H**). Consistently, similar changes were observed in the BLM-induced PF mouse model (**Figure 1I**).

### Deletion of MFAP5 Restores Elastic Fiber Homeostasis to Ameliorate PF

To examine the role of MFAP5 in the development of PF *in vivo*, we generated fibroblast-specific MFAP5 knockout mice (MFAP5^FBKO^) (**Figure 2A**). Both hematoxylin and eosin (H&E) and Masson’s trichrome staining (Masson) analyses of lung sections revealed severe fibrotic lesions in BLM-treated MFAP5^f/f^ mice, characterized by increased inflammatory infiltration, alveolar structure disruption, and collagen deposition (**Figure 2B**). Also, lung damage algorithm (LDA) analysis showed significantly reduced fibrotic areas in MFAP5^FBKO^ mice than in MFAP5^f/f^ mice challenged by BLM (**Figure 2C**). In line with histological changes, micro-computed tomography (micro-CT) analysis revealed that MFAP5^FBKO^ mice exhibited substantially reduced fibrotic regions and alleviated pulmonary consolidation relative to MFAP5^f/f^ mice following BLM treatment (**Figure 2D**). Furthermore, EFA demonstrated that the aberrant EF features observed in BLM-treated MFAP5^f/f^ mice were largely restored in MFAP5^FBKO^ mice, indicating that MFAP5 deficiency rescues EF structural characteristics during PF (**Figure 2E**). Consistently, global MFAP5 knockout (MFAP5^-/-^) mice also exhibited significantly reduced PF (**Figure 2F-G**). EFA analysis further confirmed that EF architecture in MFAP5^-/-^ mice returned toward a near-normal pattern following BLM injury (**Figure 2H-I**).

**Figure 2.**
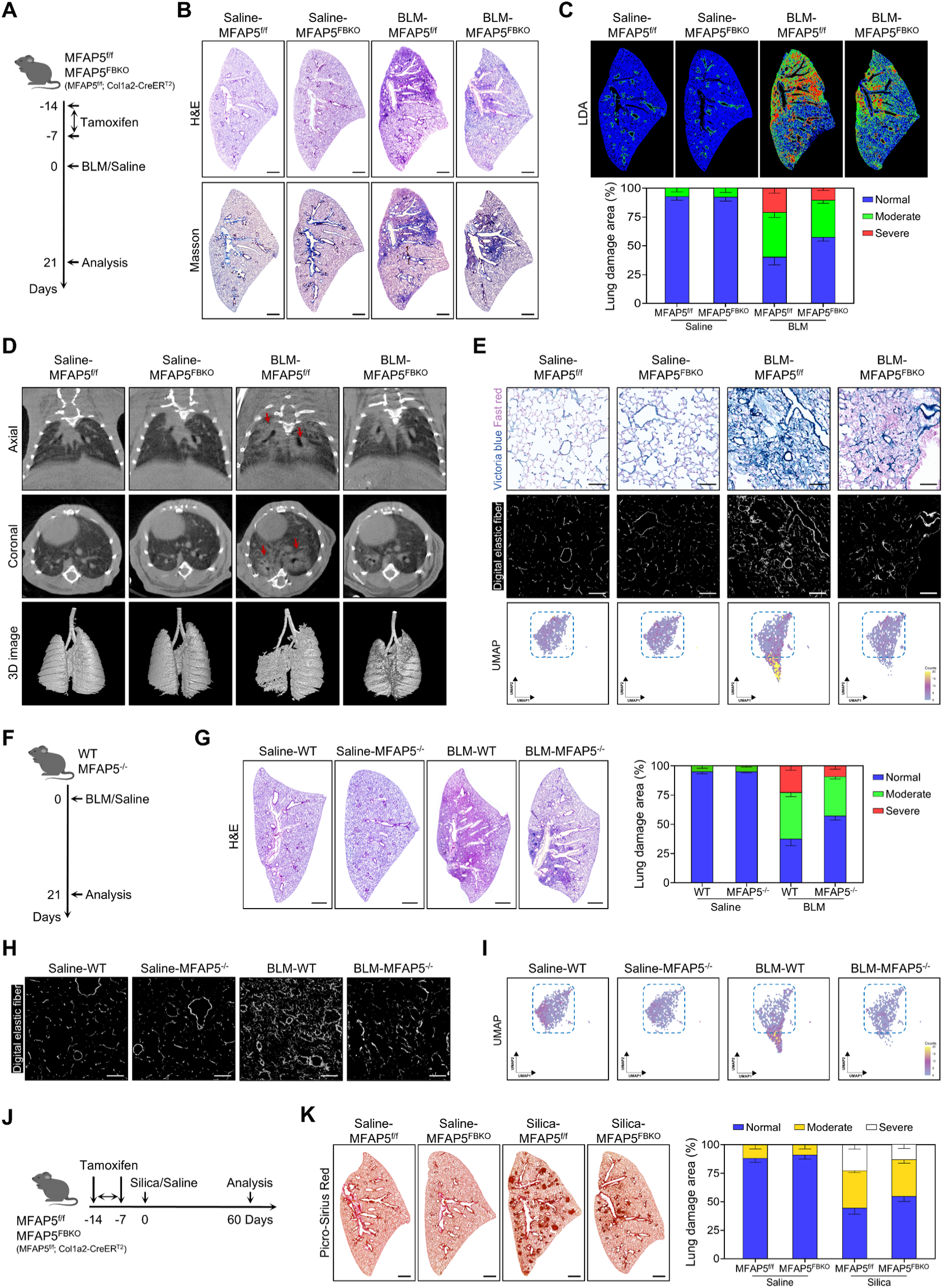
Fibroblast-specific loss of MFAP5 attenuates lung fibrosis and preserves elastic fiber homeostasis. **A**, Schematic protocol for the fibroblast-specific MFAP5 knockout (MFAP5^FBKO^) mice assays. Mice were intraperitoneally injected with 75 mg/kg tamoxifen once daily for 5 consecutive days, followed by intratracheal instillation of bleomycin (BLM) or saline. **B**, Representative hematoxylin and eosin (H&E) and Masson’s trichrome staining (Masson) images of lung sections from MFAP5^f/f^ and MFAP5^FBKO^ mice exposed to BLM or saline. Scale bars=1 mm. **C**, Representative images and quantitative results of lung damage algorithm (LDA) analysis of lungs from MFAP5^f/f^ and MFAP5^FBKO^ mice treated with BLM or saline (n=6-8). **D**, Representative micro-computed tomography (micro-CT) images of lung density. **E**, Elastic fiber algorithm (EFA) analysis showing changes in elastic fiber (EF) organization between MFAP5^f/f^ and MFAP5^FBKO^ mice following BLM challenge. Scale bars=100 μm. **F**, Schematic diagram of the experimental protocol for the global MFAP5 knockout (MFAP5^-/-^) mice subjected to BLM-induced pulmonary fibrosis (PF). **G**, Representative H&E staining images and LDA analysis of lungs from MFAP5^-/-^ mice and wild-type (WT) littermates after BLM administration (n=6-8). Scale bars=1 mm. **H**, Representative digital EF images illustrating EF distribution patterns in MFAP5^-/-^ and WT mice. **I**, UMAP visualization of EFA distribution profiles demonstrating distinct EF remodeling patterns in MFAP5-deficient versus control lungs. Scale bars=100 μm. **J**, Schematic diagram of the silica-induced PF in MFAP5^FBKO^ mice and MFAP5^f/f^ littermates. **K**, Representative Picro-Sirius Red staining images and LDA analysis showing reduced fibrosis severity in MFAP5^FBKO^ mice compared with their littermates exposed to silica (n=6-8). Scale bars=1 mm. Data represent the mean±SD. Comparisons of parameters were performed with 2-way ANOVA, followed by Tukey honestly significant difference test for multiple comparisons.

Additionally, we constructed a silica-induced PF mice model to further investigate the role of MFAP5 in a long-term PF model with chronic inflammation and irreversible fibrosis (**Figure 2J**). Sirius Red staining distinguished that silica treatment caused prominent inflammatory infiltration and extensive granuloma formation in the lungs of MFAP5^f/f^ mice. In contrast, lungs from silica-treated MFAP5^FBKO^ mice exhibited significantly reduced inflammation, granuloma size, and fibrosis severity (**Figure 2K**). Collectively, these results indicate that MFAP5 is essential for sustaining fibrotic remodeling, and its loss effectively ameliorates PF and restores EF homeostasis.

### MFAP5⁺ Fibroblasts Exhibit a Pathological Fate Bias During PF Progression

To explore the potential involvement of MFAP5 in fibroblast fate decisions during PF progression, we performed a detailed reanalysis of the scRNA-seq dataset from a BLM-induced PF mouse model (GSE207851). To better capture the temporal dynamics of fibroblast states, we redefined the groups based on the timing of BLM administration to reflect distinct pathological phases: day 0 without BLM treatment was defined as the baseline, day 1 to 7 post-BLM challenge as the inflammatory phase, day 8 to day 14 along with day 21 as the fibrotic phase, and days 28, 35, and 56 collectively as the resolving phase (**Figure 3A**). UMAP projection revealed the heterogeneity of MFAP5^+^ fibroblasts, which were further categorized into alveolar fibroblasts, adventitial fibroblasts, pathological fibroblasts, proliferative fibroblasts, and inflammatory fibroblasts based on distinct gene expression signatures (**Figure 3B-C**). Cellular proportion analysis showed that the pathological fibroblasts became dominant in the fibrotic phase and subsequently declined during the resolving phase. Similarly, proliferative fibroblasts, marked by the co-expression of pro-proliferation genes (UBE2C, BIRC4, and CCDC20) and profibrotic genes (CTHRC1, SPP1, and POSTN), exhibited a comparable temporal pattern. Moreover, consistent with the defined stages, inflammatory fibroblasts were predominantly enriched in the inflammatory phase (**Figure 3D**). Collectively, these findings underscore the temporal plasticity of MFAP5⁺ fibroblasts, which progressively transition into pathogenic states during fibrosis and partially revert during resolution. This dynamic remodeling process implicates MFAP5 as a key regulator of fibroblast activation involved in fibrosis progression.

**Figure 3.**
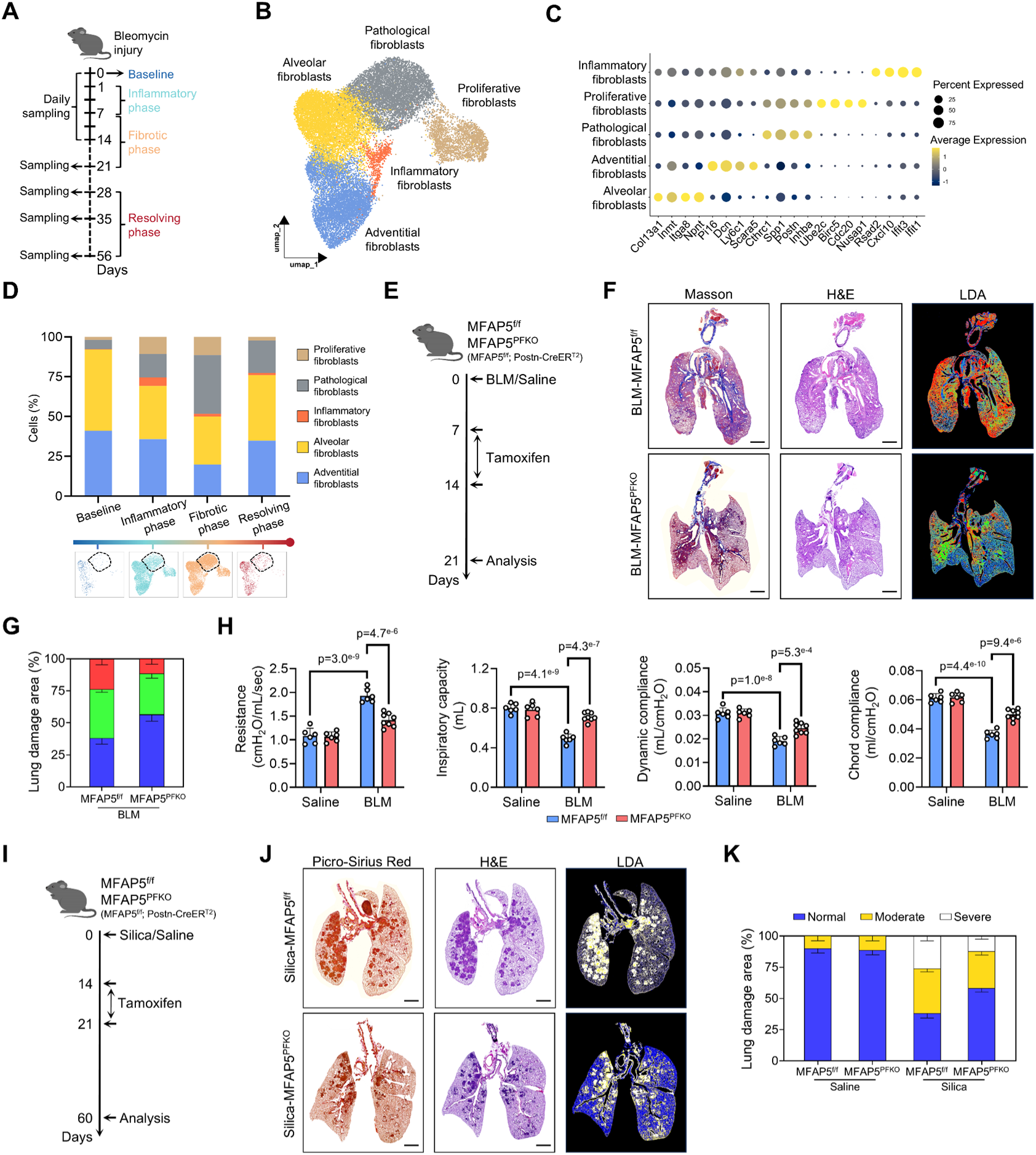
Pathological fibroblast-specific MFAP5 deletion protects against pulmonary fibrosis. **A**, Schematic overview of the experimental design and temporal grouping of single-cell RNA-seq datasets from bleomycin (BLM)-induced pulmonary fibrosis (PF) mice (GSE207851). **B**, UMAP projection of fibroblasts highlighting the heterogeneity of MFAP5⁺ fibroblast populations. **C**, Annotation of MFAP5⁺ fibroblast subpopulations, including alveolar, adventitial, pathological, proliferative, and inflammatory fibroblasts, based on distinct marker gene expression profiles. **D**, Temporal distribution of fibroblast subpopulations across disease phases. **E**, Experimental strategy for conditional deletion of MFAP5 in pathological fibroblasts (MFAP5^PFKO^) following tamoxifen administration in established BLM-induced PF models. **F**, Representative Masson’s trichrome (Masson), hematoxylin and eosin (H&E) staining, and lung damage algorithm (LDA) analysis images of lung sections from MFAP5^FBKO^ mice compared with MFAP5^f/f^ controls exposed to BLM. Scale bars=2 mm. **G**, Quantification of fibrosis severity by LDA analysis (n=6-8). **H**, Pulmonary function testing (PFT) showing the airway resistance, inspiratory capacity, dynamic compliance, and chord compliance in MFAP5^PFKO^ mice and MFAP5^f/f^ mice. **I**, Schematic protocol for the pathological fibroblast-specific MFAP5 deficiency mice assays in silica-induced PF models. **J**, Representative Picro-Sirius Red, H&E staining, and LDA analysis images of lung sections from MFAP5^FBKO^ mice compared with MFAP5^f/f^ controls exposed to silica. Scale bars=2 mm. **K**, Quantification of fibrosis severity of MFAP5^FBKO^ and MFAP5^f/f^ mice exposed to silica or saline by LDA analysis (n=6-8). Data represent the mean±SD. Comparisons of parameters were performed with 2-way ANOVA, followed by Tukey honestly significant difference test for multiple comparisons.

### Loss of MFAP5 in pathological fibroblasts reverses PF

To directly assess the contribution of MFAP5 in pathological fibroblasts, we generated pathological fibroblast-specific deficiency mice (MFAP5^PFKO^). After the establishment of PF models, tamoxifen was administered to ablate MFAP5 expression specifically within activated fibroblast populations (**Figure 3E**). Masson staining revealed a marked reduction in collagen deposition and fibrotic remodeling in MFAP5^PFKO^ lungs. Consistently, H&E staining and LDA analysis demonstrated that MFAP5^PFKO^ mice exhibited markedly alleviated fibrosis severity compared with MFAP5^f/f^ littermate controls (**Figure 3F-G**). Furthermore, pulmonary function test (PFT) revealed significant improvements in multiple respiratory parameters, including decreased airway resistance and increased inspiratory capacity, dynamic compliance, and chord compliance (**Figure 3H**). These results indicated overall recovery of lung mechanics and reduced fibrotic stiffness. Similar results were observed in the silica-induced PF model. MFAP5 deletion attenuated granulomatous lesion formation and severity (**Figure 3I-K**). In a word, these findings suggest that MFAP5 is essential for maintaining the profibrotic phenotype of pathological fibroblasts, and its loss facilitates the resolution of fibrosis even after disease onset.

### MFAP5 Drives Pulmonary Fibroblast Activation

To delineate the role of MFAP5 in regulating fibroblast states during PF progression, we performed a detailed reanalysis of the scRNA-seq dataset from IPF patients (GSE136831). MFAP5-positive (MFAP5^+^) fibroblasts were identified from the dataset and further stratified into distinct transcriptional subpopulations (**Figure 4A**). The Wnt-responsive fibroblasts (SFRP1^+^SFRP2^+^SFRP4^+^), the TGFβ/BMP-responsive fibroblasts (TGFBR3^+^SMURF2^+^HTRA3^+^), and the inflammatory fibroblasts (IL6^+^CXCL2^+^) were identified (**Figure 4B-C**). Gene Ontology (GO) enrichment analysis revealed distinct transcriptional programs and biological functions associated with each fibroblast subpopulation (**Figure 4D**). As expected, a unique enrichment of pathological (POSTN^+^CTHRC1^+^) fibroblasts in IPF patients was observed (**Figure 4E**). Together, these results suggest that MFAP5 contributes to fibroblast activation and fibrotic remodeling in PF by orchestrating fibroblast heterogeneity in response to cytokine signals.

**Figure 4.**
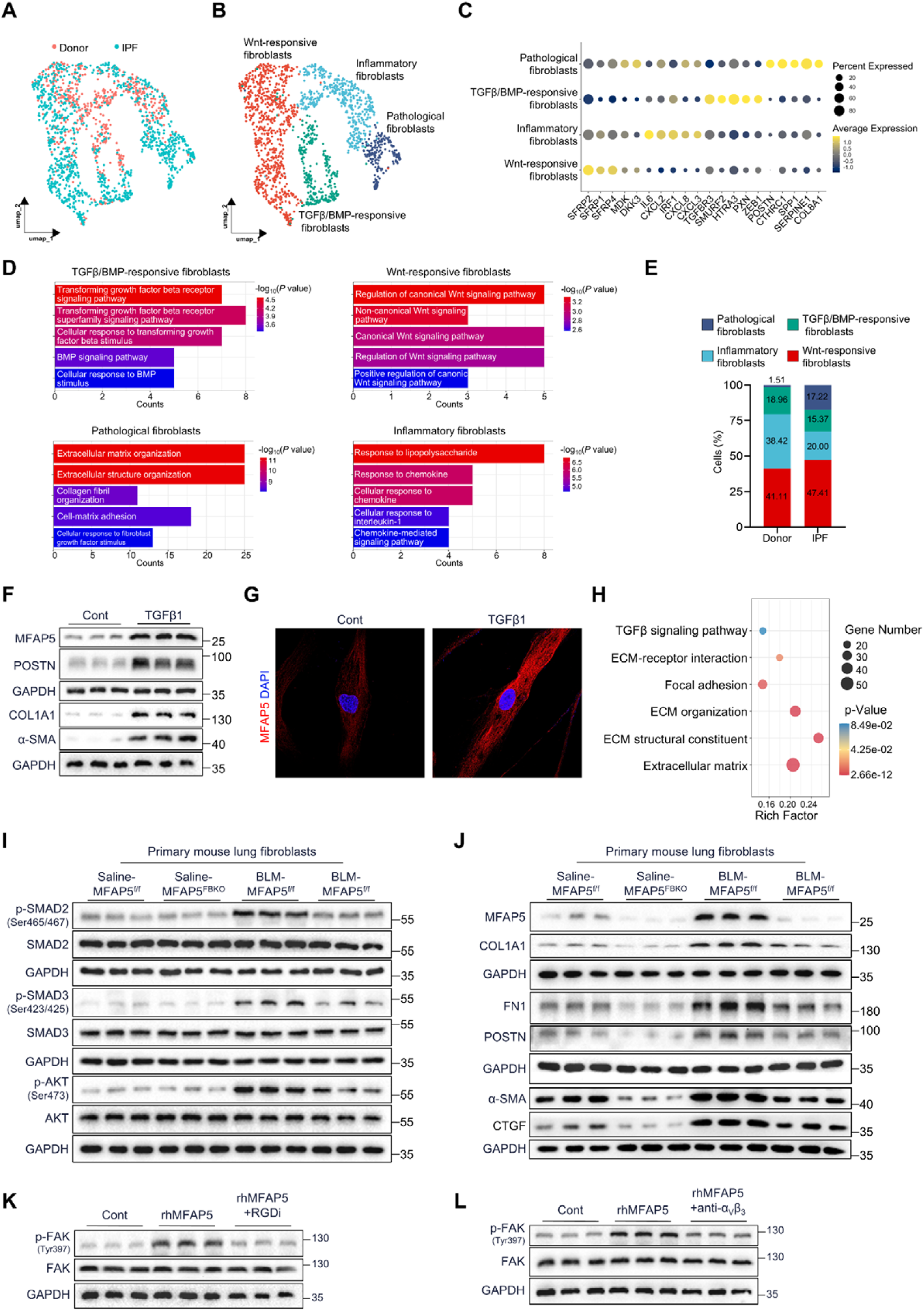
MFAP5 promotes pulmonary fibroblast activation via integrin αvβ3-FAK signaling. **A**, UMAP projection of MFAP5⁺ fibroblasts derived from the scRNA-seq dataset of idiopathic pulmonary fibrosis (IPF) patients (GSE136831). **B** and **C**, Identification of transcriptionally distinct MFAP5⁺ fibroblast subpopulations, including Wnt-responsive, TGFβ/BMP-responsive, inflammatory fibroblasts, and pathological fibroblasts. **D**, Gene Ontology (GO) enrichment analysis showing distinct biological processes and pathways associated with each subpopulation. **E**, Enrichment of MFAP5⁺ pathological fibroblasts in IPF patient lungs. **F**, Western blot analysis showing that TGFβ1 stimulation upregulates MFAP5, COL1A1, α-SMA, and POSTN in human primary pulmonary fibroblasts (HPFs). **G**, Immunofluorescent staining confirming increased MFAP5 expression in HPFs following TGFβ1 treatment. **H**, Bubble plot showing enrichment analysis of differentially expressed genes from RNA-seq of MFAP5-silenced HPFs. **I**, Western blot analysis of mouse primary pulmonary fibroblasts (MPFs) from MFAP5^FBKO^ and MFAP5^f/f^ mice showing reduced ECM proteins and fibrotic markers upon MFAP5 deletion. **J**, Reduced TGFβ signaling activation in MPFs isolated from MFAP5^FBKO^ mice compared with those from MFAP5^f/f^ controls following BLM challenge. **K** and **L**, Western blot analysis showing that recombinant human MFAP5 (rhMFAP5) induces FAK phosphorylation (p-FAK) in HPFs, which is abolished by treatment with an RGD-blocking peptide (RGDi) or an anti-integrin αvβ3 neutralizing antibody. Data represent the mean±SD. Cont, control; and TGFβ1, transforming growth factor β1.

We then investigated the role of MFAP5 in regulating fibroblast activation. TGFβ1 treatment significantly increased the protein levels of MFAP5 in human primary pulmonary fibroblasts (HPFs), accompanied by the upregulation of fibrotic markers such as COL1A1, α-SMA, and POSTN (**Figure 4F**). Immunofluorescent staining further confirmed that MFAP5 expression was upregulated by TGFβ1 treatment (**Figure 4G**). Targeting MFAP5 small-interfering RNA oligonucleotides (siMFA) were transfected into HPFs to silence its expression. Subsequent RNA sequencing (RNA-seq) analysis revealed that silencing of MFAP5 significantly downregulated genes enriched in “TGFβ signaling pathway”, “ECM-receptor interaction”, “Focal adhesion”, “ECM organization”, “ECM structural constituent”, and “Extracellular matrix” (**Figure 4H**). Western blot analysis further confirmed these findings. In mouse primary pulmonary fibroblasts (MPFs), MFAP5^FBKO^ mice exhibited decreased levels of ECM proteins (COL1A1, FN1, and POSTN), as well as canonical profibrotic markers (α-SMA and CTGF) (**Figure 4I**). These were accompanied by reduced activation of TGFβ signaling pathways compared to MFAP5^f/f^ littermate controls following BLM challenge (**Figure 4J**). Additionally, MFAP5 contains an RGD motif that binds integrins to participate in multiple disorders^5^. We found that treatment with recombinant human MFAP5 (rhMFAP5) rapidly induced FAK phosphorylation (p-FAK) in HPF. However, utilizing a competitive RGD peptide (RGDi) (**Figure 4K**) or a specific integrin αvβ3 antibody (anti-αVβ3) (**Figure 4L**) abolished rhMFAP5-induced FAK phosphorylation. Together, these results demonstrate that MFAP5 promotes profibrotic activation of human pulmonary fibroblasts by amplifying TGFβ signaling and extracellular matrix production, underscoring its functional importance in fibrotic remodeling.

## DISCUSSION

EF forms a highly complex, branching, and anisotropic meshwork to exert biological function in the lung^12^. Traditional histological and imaging approaches are difficult to capture the intrinsic network-level architecture of EF. In this study, we developed a high-dimensional EFA to address these limitations. By converting stained lung sections into digital EF-skeleton maps, over 100 architectural features that describe both local and global network characteristics were identified. The consistent shift in UMAP space in BLM-treated lungs indicates that EF remodeling during fibrosis follows a reproducible trajectory rather than occurring stochastically.

EF assembly is regulated by multiple extracellular cues and matrix-associated proteins^13^, yet the upstream factors that govern EF disorganization during fibrosis remain poorly defined. Through gene variation-rate analysis spatially, MFAP5 was identified as a curated candidate of EF-regulated gene. MFAP5 upregulation in both human PF and mouse models, along with parallel increases in established fibrotic markers, was consistent with previous reports^14^. Fibroblast-specific and global MFAP5 knockout mice demonstrated that loss of MFAP5 significantly attenuated fibrosis and restored EF architecture both in acute and chronic PF models. These data strongly support a causal role for MFAP5 in driving EF disorganization and PF progression, highlighting MFAP5 as a potential therapeutic target. Notably, studies in intrinsic dermal aging have shown that excessive MFAP5 disrupts EF assembly and induces fiber thickening and disorganization^15^. These findings support MFAP5 as a conserved regulator of EF homeostasis and a potential therapeutic target for ECM-related degenerative diseases.

Single-cell transcriptomic profiles showed that MFAP5 is predominantly expressed in fibroblasts, highlighting its cell-type-specific role within the stromal compartment. Fibroblast-ECM-integrin signaling critically drives fibrotic remodeling^16^. Activated fibroblasts sense and respond to matrix stiffness through integrin-mediated adhesion and mechanotransduction, which amplifies TGF-β signaling and promotes excessive ECM deposition^17^. The presence of an RGD motif in MFAP5 enables it to bind integrin αvβ3, involved in fracture healing^18^ and Notch signaling activation^19^. Consistently, our data demonstrated that MFAP5 promotes fibroblast activation and ECM production through integrin αvβ3-dependent signaling that amplifies the TGF-β pathway. EF assembly requires a precisely coordinated interaction among elastin, microfibrils, and associated glycoproteins. Aberrant MFAP5 expression may disrupt this balance, leading to disorganized EF architecture. Together, these findings link MFAP5-mediated fibroblast activation to EF dysregulation in PF.

While EFA provides deep structural quantification, it relies on fixed-timepoint histology. Although real-time monitoring of dynamic EF remodeling remains technically challenging, it represents an important avenue for future research. Insights gained from such studies could improve our understanding of the temporal progression of EF disorganization and inform strategies to preserve or restore EF integrity in fibrotic diseases. Gene variation-rate analysis and knockout models implicated MFAP5 as a regulator of EF homeostasis in PF, but the precise molecular interactions governing EF assembly remain to be fully dissected. Future work should examine how MFAP5 interacts with fibrillins, elastin, and crosslinkers.

In conclusion, this study provided the first quantitative, high-dimensional structural profiling of EF and identified MFAP5 as a critical regulator whose dysregulation drives EF disorganization and PF progression. Therapeutic targeting of MFAP5 may represent a novel strategy for restoring ECM integrity and alleviating fibrosis.

## METHODS

### Clinical sample collection

Lung tissues were obtained from patients with pulmonary fibrosis corresponding healthy controls enrolled in the lung transplantation programs at Shenzhen People’s Hospital and Wuxi People’s Hospital. Individuals in the control group had undergone transplant evaluation but were excluded from transplantation eligibility.

### Animal experiments design

Both male and female C57BL/6 mice were used throughout the study. All animals were purchased from Cyagen (Jiangsu, China) and maintained under specific pathogen-free conditions with a 12 h light/dark cycle at controlled room temperature. To generate MFAP5 homozygous knockout mice (MFAP5^-/-^), MFAP5 heterozygous mice (MFAP5^+/-^) were intercrossed to obtain MFAP5^-/-^ offspring and their wild-type (WT) littermate controls. For the establishment of fibroblasts (MFAP5^FBKO^) or pathological fibroblasts (MFAP5^PFKO^)-specific MFAP5 knockout mice, MFAP5 floxed mice (MFAP5^f/f^) were bred with tamoxifen-inducible fibroblasts-specific Col1a2-CreERT2 mice or pathological fibroblasts-specific Postn-CreERT2 mice. MFAP5^f/f^ Cre^+^ mice and corresponding MFAP5^f/f^ Cre^-^ littermates received tamoxifen (75 mg/kg, MedChemExpress, USA) via intraperitoneal injection once daily for 5 consecutive days to induce recombination. To establish the bleomycin-induced pulmonary fibrosis model, 8-12-week-old mice received a single intratracheal administration of bleomycin (Nippon, Japan) at 3 mg/kg, and fibrosis was allowed to develop over a 3-week period. For the silica-induced pulmonary fibrosis model, mice were given a one-time intratracheal instillation of silica (Sigma, USA) at 300 mg/kg, and fibrosis was allowed to progress for 2 months.

### Elastic Fiber Algorithm

Elastic Fiber Algorithm (EFA) was developed to digitize and quantify the abstract characteristics of elastic fibers (EF). In this workflow, Victoria blue staining was used to label EF, while Fast Red staining marked cell nuclei. An additional orthogonal was incorporated to supplement a third color vector required for subsequent color deconvolution. Whole-slide lung images were segmented into multiple independent 2000×2000-pixel tiles. A residual convolutional neural network was then utilized to remove background regions and retain tissue-only areas. Color deconvolution was performed using the Ruifrok method to isolate EF architecture from the stained images. Afterward, images were denoised and binarized to produce digital representations of thousands of individual fibers and branch points. More than 100 EF-related features, such as fiber length, width, and curvature, were subjected to principal component analysis (PCA) for dimensionality reduction, followed by UMAP visualization. In the resulting embedding, the distance between points directly reflects the differences in EF characteristics across images.

### Gene variation-rate analysis

Gene variation-rate analysis was developed to investigate the relationship between individual genes and fibrotic hotspots. Briefly, fibrotic hotspots were identified by mapping single-cell RNA sequencing data (GSE207851) onto spatial transcriptomics (S-BSST1409) using cell2location. The spatial probability of *Postn^+^Cthrc1^+^* fibroblasts was used to indicate fibrosis severity and to generate coordinate-based heatmaps and contour maps. A radius of 500 pixels around each hotspot center was defined to delineate the fibrotic niches. Gene variation-rate analysis was then performed across all fibrotic hotspots. The curves described the relationship between gene expression levels and the distance from the hotspots.

### Histological analysis of lung tissues

Lung tissue samples were fixed in 4% formaldehyde, dehydrated through a graded ethanol series, cleared in xylene, and embedded in paraffin. Sections were stained with hematoxylin and eosin (H&E), Masson’s trichrome, or Sirius Red to evaluate general tissue morphology, collagen deposition, and fibrosis. All stained sections were scanned using the Aperio VERSA 8 system (Leica, USA).

### Cell lines and cell culture

Human pulmonary fibroblasts (HPFs) were purchased from Meisen Chinese Tissue Culture Collections (Zhejiang, China). Cells between passages 3 and 8 were used for all experiments. Mouse primary pulmonary fibroblasts (MPFs) were isolated following established protocols. Briefly, lung tissue was harvested under sterile conditions, minced into small fragments, and digested enzymatically with HyqTase. The resulting cell suspension was washed, centrifuged, and plated in DMEM supplemented with 10% fetal bovine serum and antibiotics. HPFs and MPFs were maintained in fibroblast medium (2301, ScienCell, USA) and cultured at 37°C in a humidified incubator with 95% air and 5% CO₂. For TGFβ1 treatment, cells were first serum-starved for 12 h, and then incubated with 10 ng/mL TGFβ1 (R&D system, USA)under 1% oxygen conditions in a Variable Oxygen Control CO₂ incubator (Thermo Fisher Scientific, USA) for 48 h.

### Western blotting

Proteins were extracted from cells or tissue samples using RIPA lysis buffer (P0013B, Beyotime, China) supplemented with 1 mM PMSF (ST507, Beyotime). Lysates were separated by SDS-PAGE and transferred onto polyvinylidene fluoride (PVDF) membranes. Membranes were blocked and incubated overnight at 4°C with primary antibodies including MFAP5 (Invitrogen, PA5-115573, 1:1000), COL1A1 (Proteintech, 25870-1-AP, 1:1000), POSTN (Novus, NBP1-30042, 1:1000), α-SMA (Proteintech, 14395-1-AP, 1:5000), GAPDH (Proteintech, 60004-1-Ig, 1:10000), SMAD2 (Cell Signaling Technology, 5339, 1:1000), phospho-SMAD2 (Cell Signaling Technology, 3108, 1:1000), SMAD3 (Cell Signaling Technology, 9523, 1:1000); phospho-SMAD3 (Cell Signaling Technology, 9520, 1:1000), AKT Cell Signaling Technology, 9272, 1:1000), phospho-AKT (Cell Signaling Technology, 9271, 1:1000), FN1 (Invitrogen, PA5-29578, 1:2000), CTGF (Proteintech, 25474-1-AP, 1:2000), FAK (HUABIO, ET1602-25, 1:1000) and phospho-FAK (HUABIO, ET1610-34, 1:1000), followed by incubation with appropriate secondary antibodies.

### Cellular immunofluorescence staining

Cells were first fixed and permeabilized, followed by blocking to reduce nonspecific binding. Samples were then incubated overnight at 4°C with primary antibodies (MFAP5, Invitrogen, PA5-115573, 1:100). After washing, nuclei were counterstained with DAPI, and fluorescent images were captured using a laser confocal microscope.

### Recombinant human MFAP5 (rhMFAP5) protein treatment

Thr rhMFAP5 protein was purchased from Novus (4914-MG, R&D system, USA). HPFs were seeded in 6-well plates and grown to the desired confluence. Cells were then treated with 1 μg/mL rhMFAP5 or an equivalent concentration of BSA as a negative control.

### RNA sequencing and data analysis

HPFs were transfected with MFAP5 siRNA or control siRNA under TGFβ1 treatment. mRNA libraries were constructed using the mRNA-seq Lib Prep Kit (ABclonal, China) following the manufacturer’s instructions. Libraries were sequenced on an Illumina NovaSeq 6000 platform, generating 150 bp paired-end reads. Raw FASTQ reads were processed with in-house Perl scripts to remove adapters and low-quality sequences. Gene expression levels were quantified as FPKM based on gene length and read counts. Genes with a fold change≥2 and P<0.05 (Student’s t-test) were considered significantly differentially expressed. Functional enrichment analysis of differentially expressed genes was conducted using the DAVID database (https://davidbioinformatics.nih.gov/) and visualized as bubble plots in R.

### Statistical Analysis

Data were analyzed with GraphPad Prism 8.0 software and were expressed as the mean±standard deviation (mean±SD). For comparisons between two groups with a normal distribution, an unpaired 2-tailed t-test was performed. When the data did not follow a normal distribution, the non-parametric Mann-Whitney U test was applied. For data involving more than two groups, one-way ANOVA was used, followed by Tukey’s honestly significant difference test for multiple comparisons. For two variables, two-way ANOVA with Tukey’s honestly significant difference test for multiple comparisons was applied. Statistical significance was defined as P<0.05.

## Notes

### Competing Interest Statement

The authors have declared no competing interest.

